# How Social Value Guides Curiosity, Information Search, and Memory in Younger and Older Adults

**DOI:** 10.64898/2026.10.09.757930

**Authors:** Megan Vaziri, Pelin Tanberg, Ryan C. Yeung, Julia Spaniol

## Abstract

Epistemic curiosity, or the desire for new knowledge, benefits emotional, physical, and cognitive health across the lifespan. Initially believed to be solely intrinsically motivated, recent work has found that extrinsic motivators can also drive epistemic curiosity. One example of the latter is the perceived social value of information, such as popularity metrics on social media platforms (e.g., upvotes on Reddit.com). However, it remains unclear whether the influence of social value cues extends to curiosity-related behaviors, such as information search and memory, and whether these effects vary with age. The current study was designed to fill these knowledge gaps. In two experiments, younger and older adults (total *N* = 367) rated their curiosity for user-generated trivia questions that varied in social value (high or low upvote counts on Reddit.com). Upvote counts were either visible (upvote visible condition) or not visible (upvote invisible group). In Experiment 1, participants had the opportunity to reveal the answer to each trivia question, a measure of information seeking. Although social value increased curiosity, it did not influence information seeking. In Experiment 2, answer reveal was mandatory and participants received a surprise recall test for the answers. Social value again increased curiosity, particularly for younger adults, but had no influence on memory. Overall, these findings replicate social-value effects on curiosity in younger and older adults, but they provide no evidence that social value motivates information search or episodic memory. These findings highlight the limits of extrinsic influences on epistemic behaviors across the adult lifespan.

## Introduction

Imagine you are scrolling social media and come across a post that has received thousands of endorsements (e.g., likes or upvotes). Do you continue scrolling, or do you find yourself curious to learn more? In the current study, we investigate how social value cues shape epistemic curiosity and its related behaviors of information search and memory in younger and older adults.

## Aging and curiosity

Normal aging is associated with a reduction in some facets of curiosity, whereas others remain stable or even improve [1–4]. Trait-level curiosity refers to curiosity as a stable component of one’s personality [5], and typically shows negative correlations with age groups [1,4,6]. In contrast, compared to younger adults, older adults experience equivalent or greater state-level curiosity ⎯ a temporary curiosity-related motivational state [7,8].

Curiosity is associated with emotional, physical, and cognitive benefits across the lifespan [7]. Curiosity-related memory benefits [9] are of particular interest for older adults, given that episodic memory tends to decline with age [10]. Laboratory studies suggest that state curiosity during learning is positively associated with recall of curiosity-inducing information (e.g., trivia answers) [2,3,11]. Swirsky & Spaniol (2023) further found that curiosity also boosted correct recognition and reduced false recognition in older adults [12]. Hence, curiosity may serve as a protective factor against episodic memory decline in older age. Curiosity also plays a key role in guiding information-seeking behavior [13]. Information seeking involves expending resources (e.g., time, effort) to gain new knowledge, and is critical for learning, social functioning, and adaptive decision making [13,14]. Older adults exhibit a greater preference for using prior knowledge instead of exploring new information compared to younger adults [15]. However, curiosity may override this preference and encourage information seeking in older adults [8]. Given that curiosity offers several functional benefits to older adults, it is essential to understand what makes older adults curious.

## Drivers of curiosity

Epistemic curiosity is generally considered as a form of intrinsic motivation, with information gain acting as a source of reward [16]. Curiosity is intimately tied to prior knowledge [17], such that curiosity is often highest when knowledge confidence is intermediate [18] or high [19,20]. However, recent work suggests that *extrinsic* factors also influence epistemic curiosity [21]. In younger adults, curiosity has been shown to depend on perceptions of the personal or social value of the information [21–23]. For example, Dubey et al. (2021) showed participants moderately popular questions from the social media platform Reddit.com. Each question was randomly assigned a high or low upvote count to elicit the perception of high and low popularity, respectively [22]. Participants were significantly more curious about high-upvote questions than low-upvote questions, suggesting that social value, as defined by upvote count, influences epistemic curiosity. Moreover, these findings also extended to curiosity-related behaviours, such that participants engaged in greater information search for high vs. low-upvote questions.

Swirsky & Spaniol (2024) applied a version of Dubey et al.’s (2021) Reddit paradigm to compare the effects of social value on curiosity and information seeking in younger and older adults. Participants in both age groups were randomly assigned to an upvote visible condition or an upvote-invisible condition, the latter providing a baseline of curiosity and information seeking in the absence of upvotes. Visible upvotes affected curiosity in both age groups, with older adults being especially curious about trivia with high upvotes, and younger adults being especially *in*curious about trivia with low upvotes. Furthermore, information seeking tracked curiosity in both age groups. The authors interpreted the age differences in the curiosity data through the lens of socioemotional selectivity theory (SST) [24] and its behavioral hallmark, the age-related positivity effect [25]. Briefly, SST stipulates that emotion-regulation goals gain influence with age, and that this motivational shift promotes an information-processing bias in favor of positive information in older adults. The age-related positivity effect could thus explain why high upvotes—indicative of positive endorsement of information by others—were more effective at piquing curiosity in older adults than in younger adults. More generally, Swirsky and Spaniol’s (2024) observations suggest that social value influences epistemic curiosity in younger and older adults, although the nature of this influence may vary across age groups.

## The current study

As the preceding literature review has shown, there is evidence that perceived social value of information affects curiosity in younger and older adults [8,22]. This finding highlights the adaptive nature of curiosity, which helps orient us to information that other people have already vetted and endorsed. However, the degree to which social value cues influence epistemic behaviors that follow the initial assessment of curiosity, such as information seeking and long-term memory, is still unclear. Understanding the functional consequences of social value cues in older adults is of particular interest, given that exploration and memory tend to decline in normal aging [10,15]. The current study aimed to fill these gaps in current knowledge. Experiment 1 was designed to replicate Swirsky and Spaniol’s (2024) findings while addressing methodological shortcomings of their study. To foreshadow, Experiment 1 corroborated Swirsky and Spaniol’s (2024) finding of social value effects on curiosity in younger and older adults, although it did not replicate the pattern of differential sensitivity to high and low social value in the two age groups that Swirsky and Spaniol had reported. Experiment 2 built upon this empirical foundation to test whether social value would influence the fidelity of younger and older adults’ recall of answers to trivia questions.

## Experiment 1

The goal of Experiment 1 was to examine the influence of social value on curiosity and information search using a modified version of Dubey et al.’s (2021) and Swirsky & Spaniol’s (2024) Reddit paradigm, in which social value, or popularity, is operationalized as the number of upvotes (high vs. low) of trivia questions. We sought to ensure that upvote effects on curiosity and information seeking would not be due to *actual* differences in item popularity, a potential confound that was not controlled in Swirsky & Spaniol’s (2024) study. To this end, we identified a set of moderately popular trivia from the subreddit “Explain Like I’m Five” and randomly assigned them high and low upvotes. Further, following Swirsky and Spaniol’s (2024) approach, half of the participants were randomly assigned to a upvote invisible condition in which upvotes were not visible. Thus, social value effects on curiosity and information seeking would take the form of Condition x Upvote Level effects. In line with Swirsky & Spaniol (2024), we expected to find social value effects in both age groups, with high upvotes having a stronger effect in older adults, and low upvotes having a stronger effect in younger adults.

## Method

### Transparency and openness

Experimental methods, planned analyses, and hypotheses were preregistered using the OSF Preregistration template and uploaded to the Open Science Framework (OSF) Repository along with deidentified data, code for analyses, and stimuli used in experimental tasks (https://osf.io/bd7gz/). We report how we determined our sample size, any data exclusions, all manipulations, and all measures in this study. The study received approval from the Toronto Metropolitan University Research Ethics Board (protocol number: 2022-237). Data collection took place from January 15, 2025, to February 21, 2025. Written informed consent was obtained digitally.

### Participants

Participants from Canada and the United States were recruited via CloudResearch’s Amazon Mechanical Turk (MTurk) Toolbox. Following Swirsky & Spaniol (2024), a target sample size of 180 was determined to slightly exceed the minimum sample size estimated via an a priori power analysis conducted using MorePower 6.0.4 [26]. One hundred sixty-eight participants would be required to achieve a power of 0.90 to detect a medium-sized (η_p_ ^2^ = .06; Swirsky & Spaniol, 2024) 3-way interaction of age group, condition (upvote visible vs. upvote invisible), and upvote level (high vs. low), assuming an α error probability of .05. To account for exclusions and potential dropouts, 214 participants were recruited. Participants who demonstrated no variability in their ratings, completed the three ratings in less than three seconds on one or more trials, indicating rushed responding, revealed no answers, completed the study multiple times, terminated the study prior to completion/had technical difficulties submitting their responses, and completed fewer than 12 years of education were excluded and replaced (n=31). Therefore, a total of *N* = 183 participants were retained, including 92 younger adults (ranging 18-35 years; 46 females; see Table 1 for distribution of ethnicities) and 91 older adults (age range: 60-93 years; 45 females; see Table 1 for distribution of ethnicities). An overview of sample characteristics is reported in Table 2.

**Table 1.** Distribution of Ethnicities in Experiment 1.

| Ethnicity | Younger Adults ($N=92$) | Older Adults ($N=91$) |
| --- | --- | --- |
|  | n(%) | n (%) |
| White | 57(61.96) | 82(90.11) |
| White/Latin American | 4(4.35) | 1(1.10) |
| White/Latin American/Indigenous |  | 1(1.10) |
| White/Black or African American | 3(3.26) |  |
| White/Indigenous | 3(3.26) |  |
| White/Chinese | 1(1.09) |  |
| White/South Asian | 1(1.09) |  |
| White/Bohemian |  | 1(1.10) |
| Black or African American | 9(9.78) | 5(5.49) |
| Black or African American/Latin American | 1(1.09) |  |
| Latin American | 5(5.43) |  |
| Arab | 1(1.09) |  |
| Filipino | 1(1.09) |  |
| Filipino/Southeast Asian | 1(1.09) |  |
| Southeast Asian | 1(1.09) |  |
| Korean | 1(1.09) |  |
| Chinese | 3(3.26) |  |
| Prefer not to answer |  | 1(1.10) |

**Table 2.**
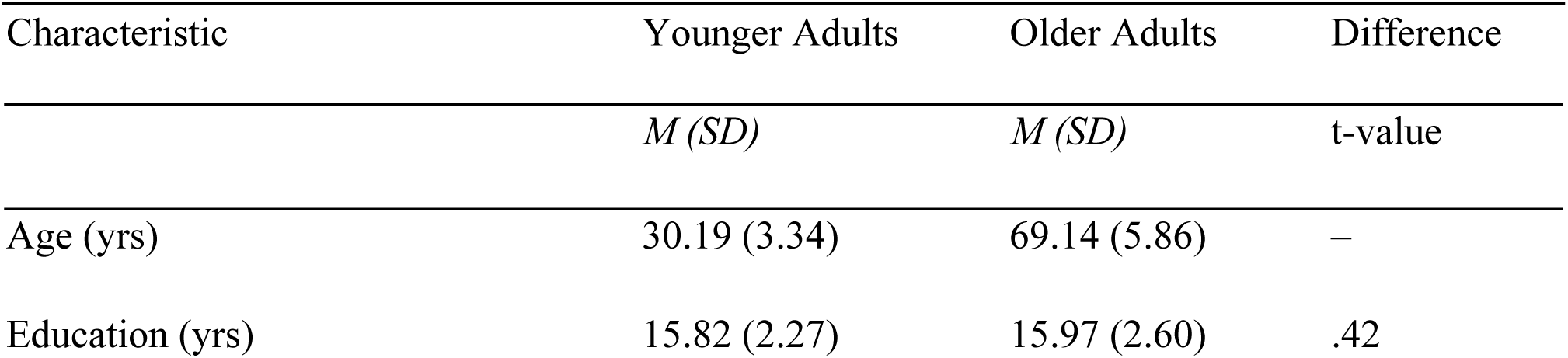
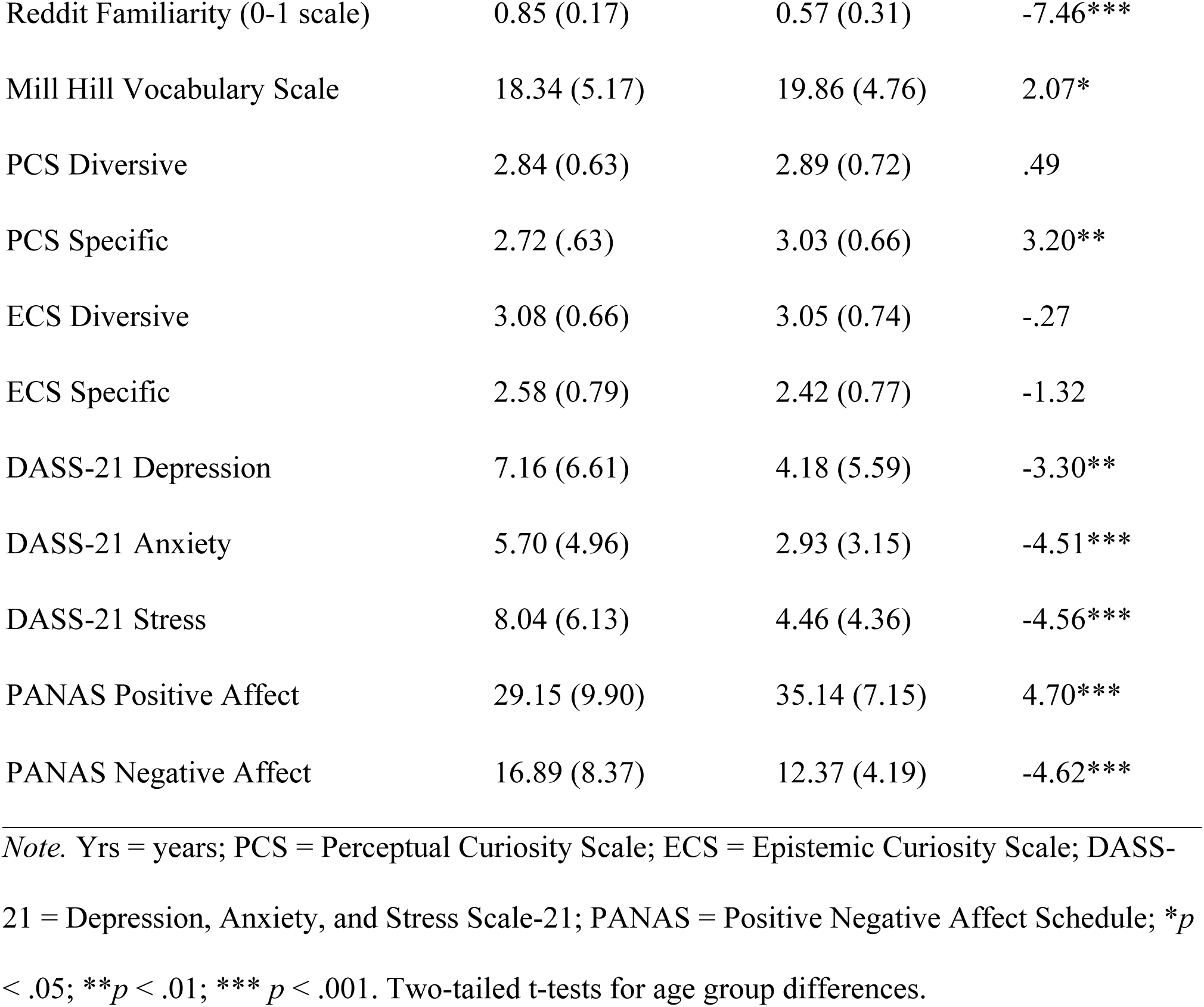
Sample Characteristics and Age Effects in Experiment 1.

### Materials

Materials consisted of 20 trivia questions and answers originally sourced from the subreddit “Explain Like I’m Five” on Reddit.com, where they had received between 200 and 600 upvotes, rendering them moderate in popularity [22]. Four additional questions were used as practice questions. Question topics included general science, medical/biological, and societal/psychological. For each question, we selected the user-generated answer with the highest upvote count. Answers were shortened as needed to conform to a 100-200-word length limit and contain 3-7 key informational elements. Original answers were edited for grammar and clarity if necessary.

In the upvote visible condition, each question was randomly assigned a low (4-95) or high (26,600-79,200) upvote level, such that there were 10 questions per upvote-level condition. Low- and high-upvote level values were randomly drawn from the upvote ranges used in Swirsky & Spaniol (2024; low: 4-99, high: 26,100-80,900). As a result, there was no systematic relationship between the actual popularity of a question and the upvote level that was shown to participants. The upvote level assigned to each question was held constant across participants to facilitate comparisons across upvote visible and upvote invisible conditions at the item level. In the upvote invisible condition, no upvotes were shown.

Participants completed a series of questionnaires indicating their age (years), gender (options: man, woman, prefer to specify, prefer not to answer), ethnicity (options: White, South Asian, Chinese, Black/African American, Filipino, Arab, Latin American, Southeast Asian, West Asian, Korean, Japanese, prefer to specify, prefer not to answer), Indigenous status, education (years), high school degree completion, English language proficiency (native language status; comfort reading/writing in English), and familiarity with Reddit.com [measured on a continuous visual analogue scale (i.e., slider) ranging from 0 to 1, where 0 was anchored as “Not at all” and 1 as “Extremely,” with participants able to choose any point along the continuum]. They also completed a battery of measures assessing trait curiosity via the Perceptual Curiosity Scale [27] and Epistemic Curiosity Scale [28], verbal intelligence using the Mill Hill Vocabulary Scale [29], and current mood states with the Positive and Negative Affect Schedule [30] and the Depression, Anxiety, and Stress Scales [31]. All tasks were programmed in PsychoPy v2024.1.4 [32] and hosted on Pavlovia (Open Science Tools, Nottingham, UK).

### Design and procedure

The study used a 2 (age group: younger adults, older adults) x 2 (condition: upvote invisible, upvote visible) x 2 (upvote level: high, low) mixed design with repeated measures on the last factor. Depending on the survey link they selected on CloudResearch’s MTurk (Experiment 1) or Prolific (Experiment 2), participants either completed the upvote visible (Experiment 1: 92 younger adults, 91 older adults; Experiment 2: 92 younger adults, 92 older adults) or upvote invisible study (Experiment 1: 92 younger adults, 92 older adults; Experiment 2: 92 younger adults, 92 older adults). Survey names, links, and advertisements on the platforms were identical.

In the upvote visible condition, participants saw the number of upvotes associated with each question, signifying the number of Reddit.com users who had endorsed the question. Participants were instructed that each question was viewed by the same number of online users and that upvote counts signified endorsement rather than variations in audience size. Before proceeding with the task, participants completed a multiple-choice question to confirm that they understood the meaning of an upvote. Participants were also instructed that upvotes were given by individuals in their own age group. In the upvote invisible condition, participants did not see upvotes.

Participants completed four practice trials to ensure they understood the task. On each trial, participants saw the questions (two high-upvote questions, two low-upvote questions) in a random order before responding to a series of prompts. The prompts required participants to rate their curiosity and their knowledge confidence [measured on a continuous visual analogue scale ranging from 0 to 1, where 0 was anchored as “Not at all” and 1 as “Extremely,” with participants able to choose any point along the continuum]. As a manipulation check, participants also rated how popular they perceived each question to be on the same scale. The manipulation check served to verify that upvote levels (high vs. low) in the upvote visible condition influenced participants’ perceptions of popularity. After completing the practice trials, participants saw the 20 questions (10 high-upvote questions, 10 low-upvote questions) in a random order and responded to the same prompts.

In the second phase of the experiment, participants were presented with each of the 20 questions in a random order. Participants in the upvote visible condition again saw the upvote level associated with each question, whereas those in the upvote invisible condition did not. For each question, the participant had the option to reveal the answer before moving to the next question. Answer reveal decisions served as a measure of information seeking. After the experiment, participants completed the demographic questionnaire and background measures (see Fig 1 for sequence of tasks), and were debriefed about the mild deception regarding upvote levels shown to participants in the upvote visible condition.

**Fig 1.**
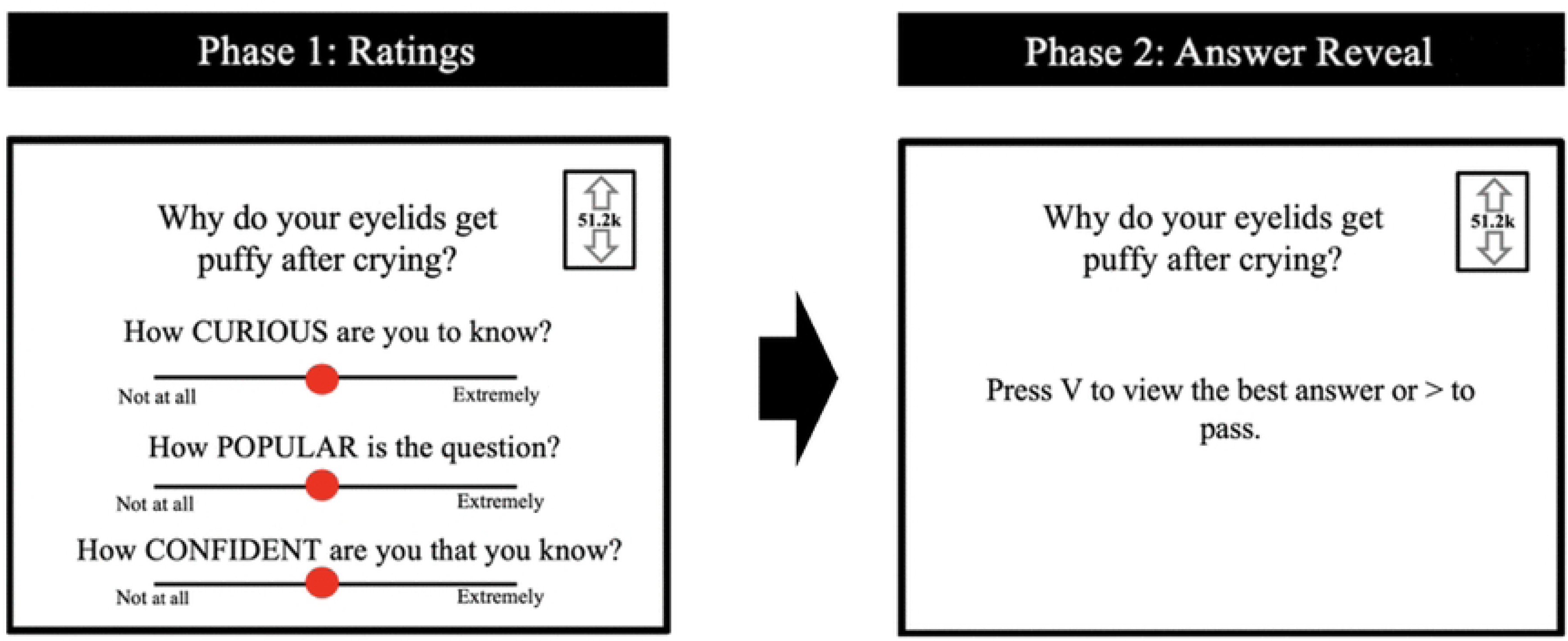
Experiment 1 Procedure. Study sequence in the upvote visible condition. The upvote invisible condition followed the same sequence, except upvote levels were not shown. In Phase 1, the rating scales were presented one at a time, with the subsequent scale appearing only after a response was made for the preceding scale. Upon finishing Phase 2, participants completed background measures and demographic questionnaires, which are not depicted in the figure.

### Analyses

All statistical analyses were carried out in R version 4.6.1 (R Core Team, 2026). Analyses followed Swirsky & Spaniol (2024)’s approach of estimating mixed-effects models with random intercepts for both participant and question identifiers to account for repeated observations within individuals and trivia items. The structure of the random effects models followed the recommendations of [33]. We analyzed popularity ratings to assess whether the manipulation of upvote levels in the upvote visible condition was successful. Predictors included age group, condition, upvote level, and interaction terms. Furthermore, we examined trial-level curiosity ratings. Predictors included age group, condition, upvote level, and interaction terms. Lastly, we analyzed information-seeking behaviour as defined by answer reveal decision (yes or no). Predictors included age group, condition, upvote level, and interaction terms.

We included random slopes for all predictors that varied within participants (i.e., upvote level) and within questions (i.e., age, condition), but only if they improved model fit. If the model was singular, we continued to reduce the model complexity to report the best-fitting maximal model, and we state the random effects structure used for that model. Generalized linear mixed models were used for continuous outcomes, using the *lmer* function, while logistic mixed models were used for binary outcomes, using the *glmer* function [34]. All models were fitted using the *bobyqa* optimizer to improve convergence and ensure more stable estimation of model parameters, particularly given the complexity of the random effects structure. In addition, for easier interpretation, odds ratio statistics for the interpretation of effect sizes (OR) [35,36], along with model-based 95% confidence intervals (CIs), are reported for all logistic regression models.

Fixed effects included age (younger, older), condition (upvote invisible, upvote visible), and upvote level (low, high). Once model fit was established, we evaluated the significance of fixed effects and their interactions using Wald chi-squared tests via the *Anova* function from the *car* package [37]. Significant fixed effects and their interactions were followed up by pairwise comparisons using the *emmeans* package [38], which computes estimated marginal means (i.e., least-squares means) based on the fitted models. Pairwise comparisons of categorical predictors were conducted using the *emmeans* function, with the Tukey adjustment applied to control for Type I error given multiple comparisons.

As per the preregistration, models initially included all variables that showed significant age differences as covariates, and these results are reported in the Supplementary Materials. However, the covariates were removed from the main analysis models for simplification. Cluster-mean-centered confidence and curiosity were also not included as predictors in these models. Despite being important variables to curiosity-related outcomes, incorporating these variables, particularly as continuous predictors involved in interaction terms with experimental factors, substantially increased model complexity. As demonstrated in preliminary analyses, models including these predictors, especially in combination with higher-order interactions and random effects, produced unstable or non-interpretable estimates despite convergence. Given these concerns, and the absence of specific preregistered hypotheses regarding their role as predictors in the information search models, confidence and curiosity were excluded to preserve model stability, interpretability, and parsimony. Additional analyses incorporating these predictors can be found in the Supplementary Materials.

## Results

### Manipulation check: Popularity ratings

To confirm the effectiveness of the social value manipulation, we regressed trial-level popularity ratings on age group, condition, and upvote level. Briefly, these analyses confirmed that the upvote levels influenced perceptions of popularity in both age groups (see Supplementary Materials).

### Curiosity ratings

We regressed trial-level curiosity ratings on age, condition, and upvote levels. The final model that converged included a random intercept and slope for upvote by participant, as well as a random by-question intercept, with correlations between the participant-level random intercept and slope estimated. There was a significant Condition x Upvote Level interaction, *χ^2^* (1) = 15.24, *p* < .001 (Fig 2). In the upvote visible condition, participants were significantly more curious about high-upvote (*M* = .57, SE = .03) than low-upvote (*M* = .50, SE = .03) questions, *b_estimate_* = .07, SE = .03, 95% CI [.00, .12], *p* = .048. In contrast, in the upvote invisible condition, there was no significant difference in curiosity for high-upvote (*M* = .56, SE = .03) and low- upvote questions (*M* = .57, SE = .03), *b_estimate_* = -.01, SE = .03, 95% CI [-.08, .05], *p* = .733. There were no significant effects involving age, all *p*s > .05. In summary, when participants saw upvotes (i.e., in the upvote visible condition but not in the upvote invisible condition), these social-value cues had a significant impact on curiosity, and this effect was similar for younger and older adults.

**Fig 2.**
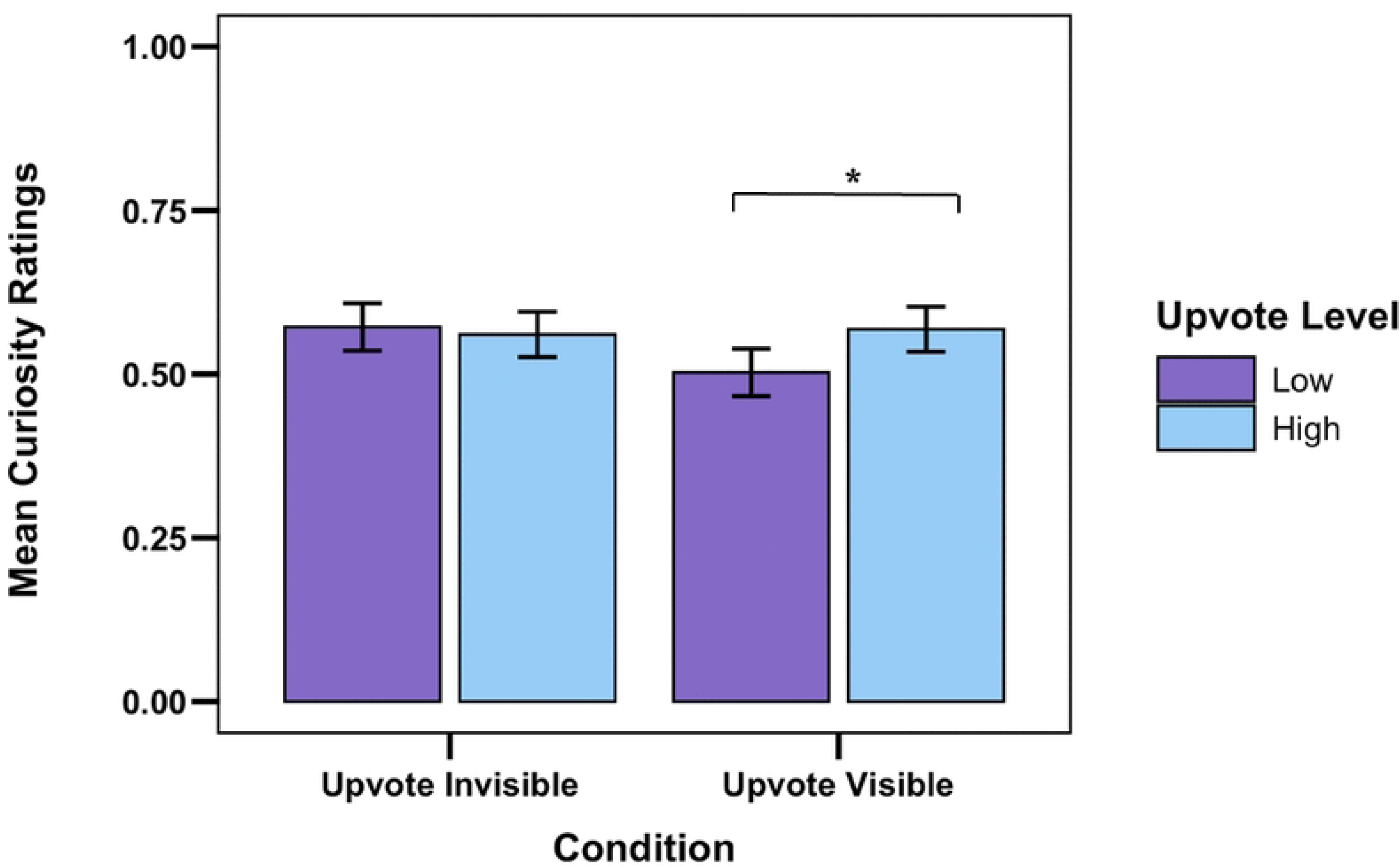
Curiosity Ratings by Condition and Upvote Level in Experiment 1. *\*p <* .05. Error bars represent one standard error above and below the estimated marginal mean.

### Information seeking

To explore potential effects of social value cues on information seeking, we regressed answer-reveal decisions on age group, condition, and upvote level. The final model that converged included a random by-participant and by-item intercept, excluding all correlations among random effects. The model did not reveal any significant main effects or interactions (all *ps* > .05). This suggests that information seeking was not significantly influenced by social value in either age group. We discuss these findings in the General Discussion.

## Experiment 2

Experiment 2 was similar to Experiment 1, but its goal was to test the effect of social value on memory. Therefore, rather than allowing participants to choose whether to reveal trivia answers, answer reveal was mandatory. At the end of the experiment, participants received a surprise memory test for the trivia answers. In line with the Experiment 1 results and prior work [8], we hypothesized that in the upvote visible condition, curiosity and memory would be higher for high- vs. low-upvote trivia. We also sought to explore the effects of social value on memory in younger and older adults. Memory performance on the trivia answer recall was quantified using semantic textual similarity [39,40] rather than traditional manual coding. Semantic similarity scores as derived from transformer-based language models (e.g., BERT) [41] are sensitive to meaning rather than exact wording [42], making them particularly well-suited for naturalistic recall tasks in which participants may accurately remember information using paraphrased language. Using semantic similarity also improved scalability and reproducibility by eliminating the need for multiple human raters and extensive inter-rater reliability procedures [41,43].

## Method

### Transparency and openness

Experimental methods, planned analyses, and hypotheses were preregistered using the OSF Preregistration template and uploaded to the OSF repository along with deidentified data, code for analyses, and stimuli used in experimental tasks (https://osf.io/wzdut/). We report how we determined our sample size, any data exclusions, all manipulations, and all measures in this study. The study received approval from the Toronto Metropolitan University Research Ethics Board (project title: Determinants of Curiosity; protocol number: 2022-237). Data collection took place from April 10, 2025, to May 21, 2025. Written informed consent was obtained digitally.

### Participants

Adult participants from Canada, the United States, the United Kingdom, and Ireland were recruited via Prolific. Target sample size and power analysis results were identical to Experiment 1. To account for exclusions and potential dropouts, 224 participants were recruited. To screen for repeat participants from Experiment 1, Prolific participants were asked to indicate whether they were also MTurk workers, and if so, to provide their MTurk Worker ID. Participants who demonstrated no variability in their ratings, completed ratings in less than three seconds on a single trial, indicating rushed responding, terminated the study prior to completion/had technical difficulties with their submission, reported ages other than 18-35 or 60+, had less than 12 years of education, and remembered no answers were excluded and replaced (n=40). Therefore, a total of 184 participants were retained, including 92 younger adults (ranging 18-35 years; 46 females; see Table 3 for distribution of ethnicities) and 92 older adults (ranging 60-99 years; 46 females; 1.09% Indigenous; see Table 3 for distribution of ethnicities). A summary of sample characteristics is reported in Table 4.

**Table 3.**
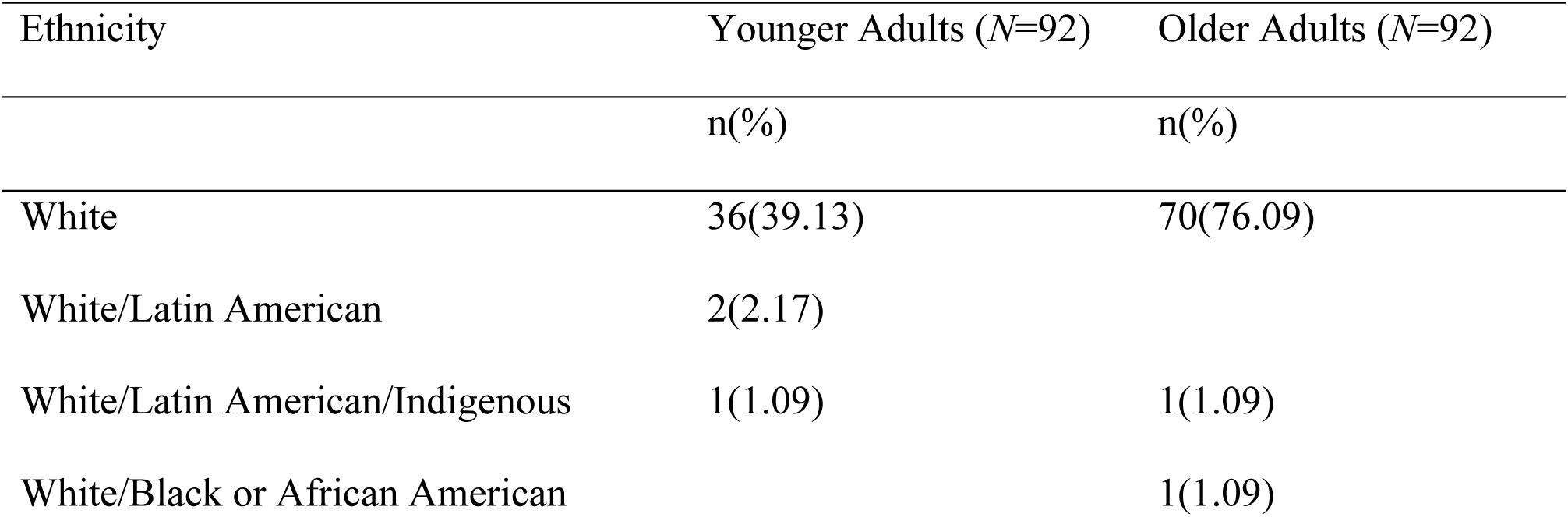
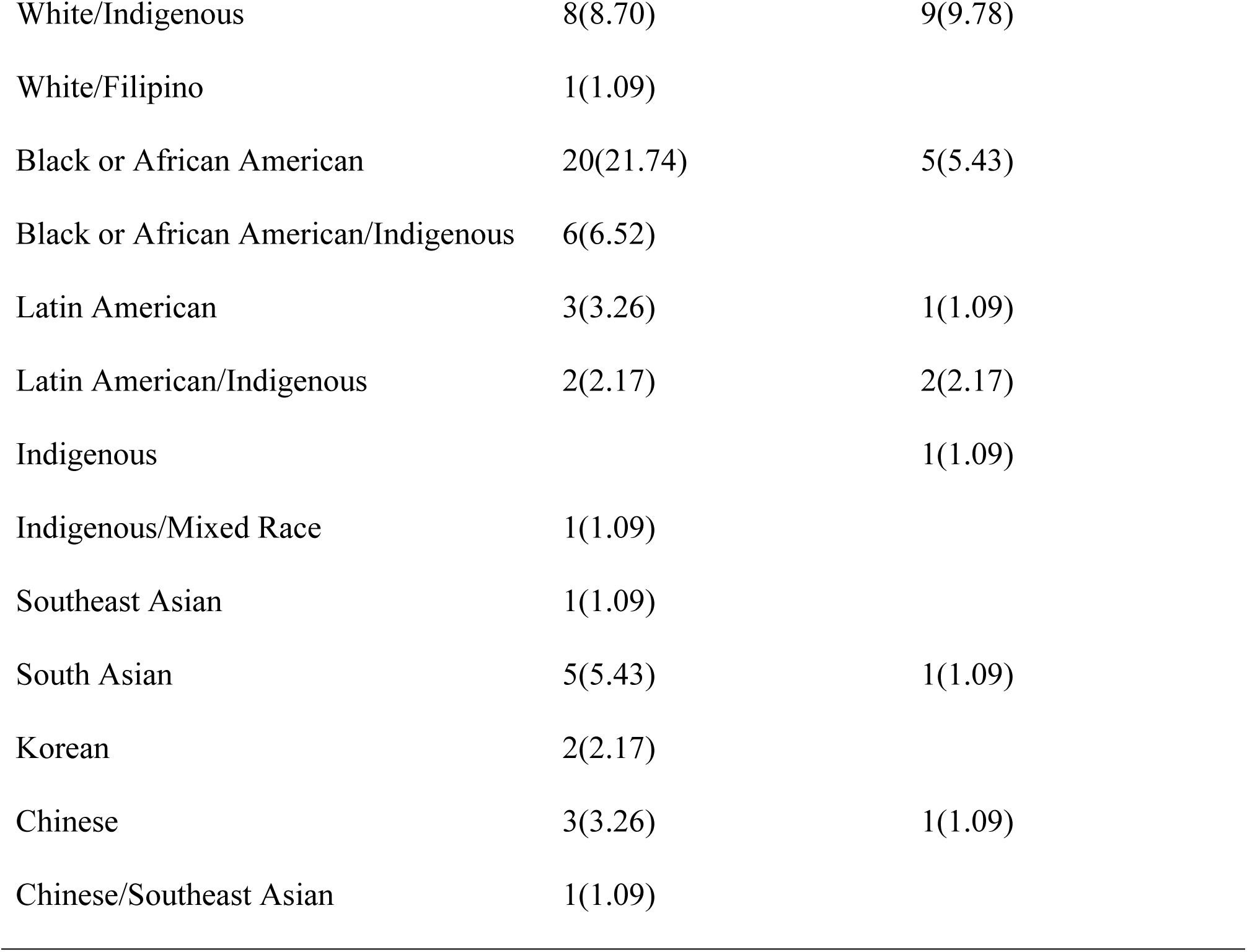
Distribution of Ethnicities in Experiment 2.

| Ethnicity | Younger Adults ($N=92$) | Older Adults ($N=92$) |
| --- | --- | --- |
| | $n(\%)$ | $n(\%)$ |
| White | 36(39.13) | 70(76.09) |
| White/Latin American | 2(2.17) |  |
| White/Latin American/Indigenous | 1(1.09) | 1(1.09) |
| White/Black or African American |  | 1(1.09) |
| White/Indigenous | 8(8.70) | 9(9.78) |
| White/Filipino | 1(1.09) |  |
| Black or African American | 20(21.74) | 5(5.43) |
| Black or African American/Indigenous | 6(6.52) |  |
| Latin American | 3(3.26) | 1(1.09) |
| Latin American/Indigenous | 2(2.17) | 2(2.17) |
| Indigenous |  | 1(1.09) |
| Indigenous/Mixed Race | 1(1.09) |  |
| Southeast Asian | 1(1.09) |  |
| South Asian | 5(5.43) | 1(1.09) |
| Korean | 2(2.17) |  |
| Chinese | 3(3.26) | 1(1.09) |
| Chinese/Southeast Asian | 1(1.09) |  |

**Table 4.**
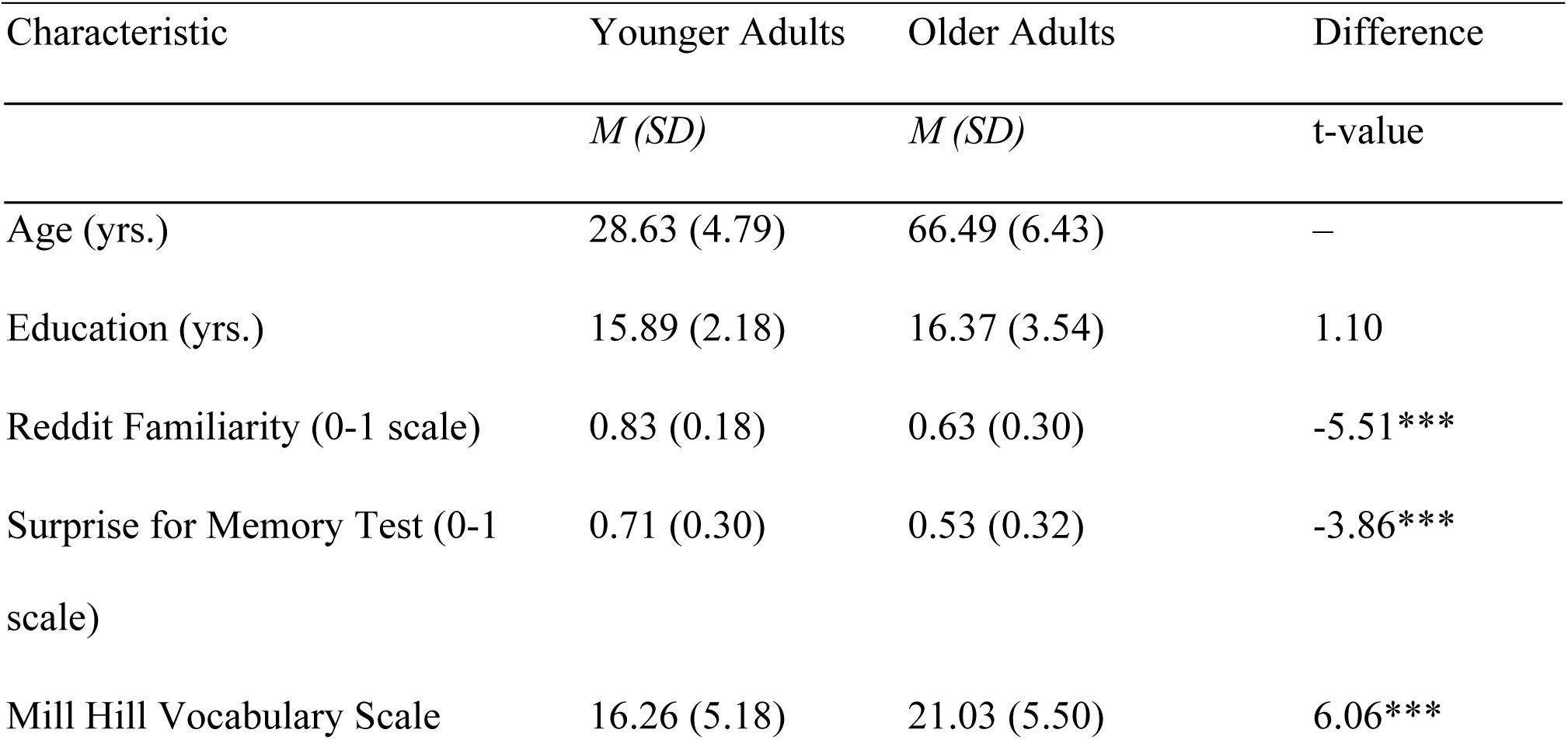
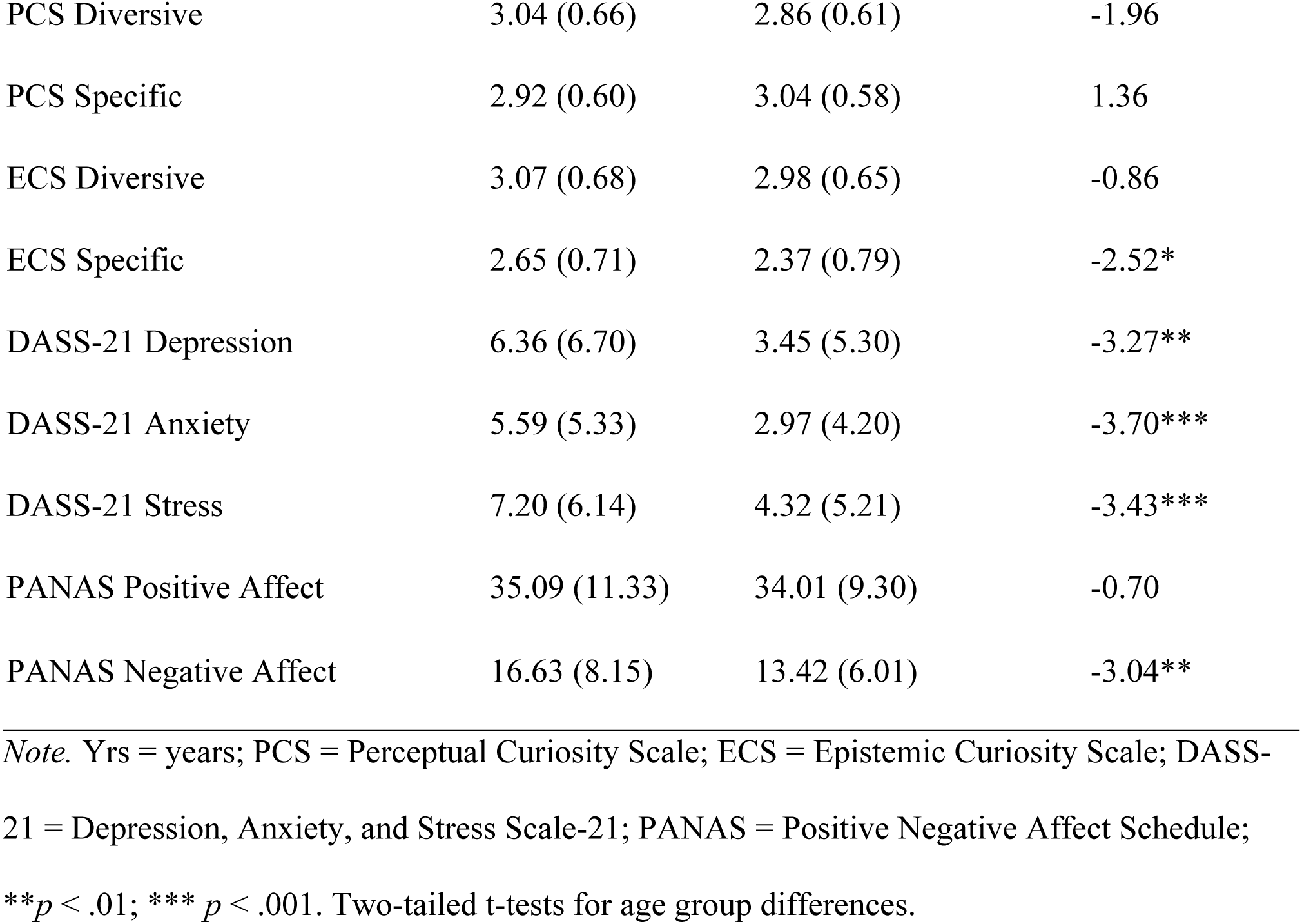
Sample Characteristics and Age Effects in Experiment 2.

### Materials, design, and procedure

Materials were identical to those used in Experiment 1. The experimental design differed from Experiment 1 in that we investigated trivia recall performance instead of information seeking. After the ratings phase, participants were presented with the 20 questions along with their answers in a random order for a minimum of 20 seconds per pair. To promote engagement, participants were instructed that after reading each pair, the question and answer would disappear, and they would have to rate their level of surprise about the answer on a continuous visual analogue scale ranging from 0 to 1, where 0 was anchored as “Not at all” and 1 as “Extremely.” Participants were able to choose any point along the continuum. They then submitted their rating using a “Submit” button before the next pair appeared. Participants in the upvote visible condition saw the upvote counts associated with each question, whereas those in the upvote invisible condition did not.

After reading each answer, participants completed an unrelated go/no-go task for one minute before beginning the third phase. The cognitive task involved digits 1-9 being presented one at a time in the centre of the screen. Numbers were presented in a random order and remained on the screen for one second each before the subsequent number was presented. Participants were instructed to press the spacebar each time three odd numbers appeared consecutively.

In the final, self-paced phase of the experiment, participants were presented with each of the 20 questions again and were asked to type as many details as they remembered about each answer into an open text box. They were instructed to press an “I don’t know” button if they did not remember any details about the answer. Participants were required to either submit their response using a “Submit” button or select the “I don’t know" button to proceed to the next question. After completing the task, participants were asked to report their surprise about receiving the memory test on a scale [continuous visual analogue scale ranging from 0 to 1, where 0 was anchored as “Not at all surprised” and 1 as “Extremely surprised,” with participants able to choose any point along the continuum]. Lastly, participants completed the demographic questionnaire items and background measures (see Fig 3 for the sequence of tasks).

**Fig 3.**
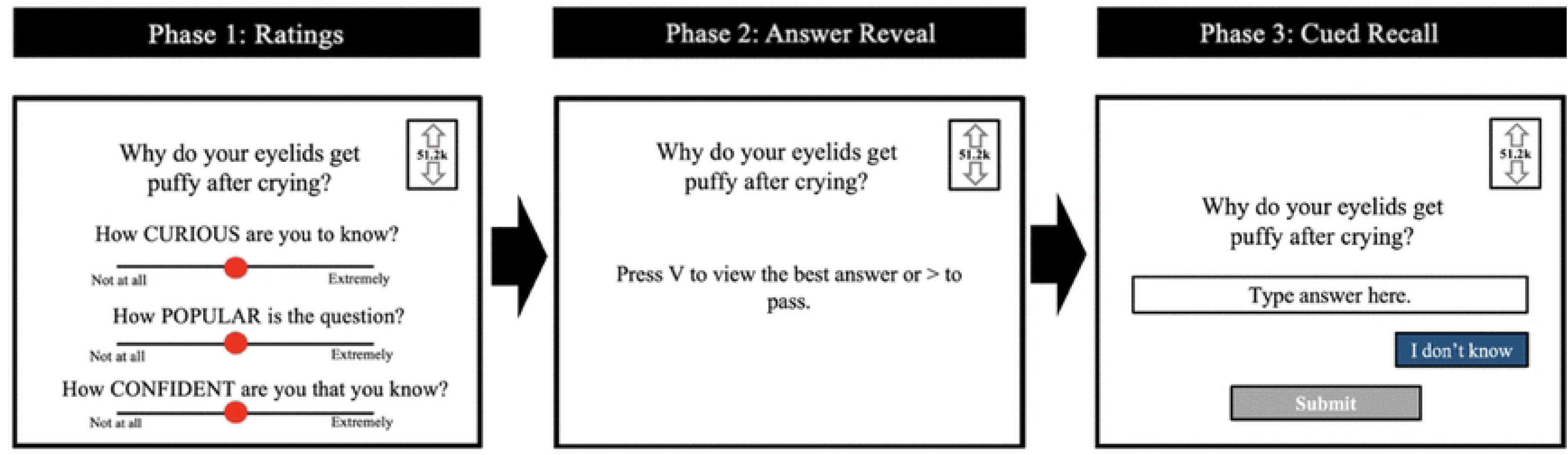
**Experiment 2 Procedure**. The figure depicts the task sequence for the upvote visible condition. The upvote invisible condition followed the same order, but upvote levels were not visible. In Phase 1, the rating scales were presented individually, with each following scale appearing only after a response was made for the preceding items. Unlike in Experiment 1, participants did not have the option to reveal the answers to the questions in Phase 2. Upon finishing Phase 3, participants completed background measures and demographic questionnaires, which are not depicted in the figure.

### Analyses

Analyses of popularity and curiosity ratings paralleled those conducted in Experiment 1. Additionally, we examined recall performance as operationalized by semantic similarity between open text recall responses and their original trivia answers (see Table 5). Predictors included age group, condition, upvote level, and their interaction terms.

**Table 5.** Sample High- and Low-Similarity Recall Passages for a Trivia Question.

| Question | Original Answer | Low-Similarity<br>Answers | High-Similarity<br>Answers |
| --- | --- | --- | --- |
| What exactly is happening when we blur our eyes on command? | Our eyes can't focus on really near things and really far things at the same time, but we can actively switch between focusing on near things (making far things look blurry) or focusing on far things (making near things look blurry). This is the process called "accommodation." | It is something to do with what is called "accommodation" between close and distance objects. (Cosine similarity score = .26)<br>We are controlling a muscle. (Cosine similarity score = .31)<br>Muscle control. (Cosine similarity score = .22) | When we blur our eyes on command, a muscle in our eyes control this movement. (Cosine similarity score = .82)<br>When we're blurring our eyes on command, it's because they're flexing a certain muscle in our eyes that causes us to lose focus and create |
|  | There is a biological lens inside each of our eyes, and when the tiny muscles connected to the lens pull on it or relax, it changes its shape, which changes the way light is focused in the eye. When you “blur your eyes on command,” you’re actually controlling those tiny muscles connected to the lens. |  | blurred vision. (Cosine similarity score = .80)<br>When we blur our eyes on command, we are controlling some muscle in our eyes that controls focus. (Cosine similarity score = .86) |

### Semantic similarity

To index participants’ memory for answers, semantic similarity scores were calculated between each cued recall response (Phase 3) and its original answer (Phase 2) using natural language processing. Briefly, pretrained transformer-based language models can be used to represent a given text as an embedding (i.e., a vector), which captures semantic relationships between texts [42]. For instance, two very semantically similar texts (e.g., “the big dog” vs. “the large canine”) would produce vectors that are very close to each other. In contrast, two very semantically dissimilar texts (e.g., “the small cat” vs. “it’s raining today”) would produce vectors that are very distant from each other. As such, higher similarity between a participant’s cued recall response and its original answer would indicate better memory for the answer since the content of their recall is highly semantically similar to the original to-be-encoded content.

Following Fowler et al. (2024), here we use the ‘multi-qa-distilbert-cos-v1’ model released by Sentence Transformers [44]. This model is based on the pretrained ‘distilbert-base- uncased’ model [45] and additionally finetuned on 215 million question-answer pairs to optimize for semantic search. Using this model, all cued recall responses and original answers were converted into embeddings (vectors containing 768 numbers), and the distance between each cued recall response and its original answer was computed in terms of the embeddings’ cosine similarity. Notably, this model produces normalized embeddings (length = 1), so cosine similarity and dot product similarity metrics are numerically identical. Cued recall responses of “I don’t know” were considered the same as “I don’t know” button responses. Otherwise, no preprocessing was applied to either the cued recall responses or the original answers. For illustrative purposes, examples of high and low similarity recall passages for one of the trivia answers are shown in Table 5.

## Results

### Manipulation check: Popularity ratings

Popularity ratings were sensitive to upvotes in the upvote visible condition (see Supplementary Materials for details), confirming the effectiveness of the social value manipulation for both age groups.

### Curiosity ratings

We regressed trial-level curiosity ratings on age, condition, and upvote level. The final model that converged included a random intercept and slope for upvote by participant, as well as a random by-question intercept, with correlations between the participant-level random intercept and slope estimated. We found a significant main effect of age, *χ^2^* (1) = 9.07, *p* = .003, which was qualified by significant two-way interactions of Age x Upvote Level, *χ^2^* (1) = 4.45, *p* = .035, and Condition x Upvote Level, *χ^2^* (1) = 34.92, *p* < .001. Finally, there was a significant Age x Condition x Upvote Level interaction, *χ^2^* (1) = 4.23, *p* = .040 (Fig 4). Separated by Age and Upvote level, younger adults were significantly less curious about low-upvote items in the upvote visible (*M* = .53, SE = .03) than upvote invisible (*M* = .67, SE = .03) condition, *b_estimate_* = .14, SE = .04, 95% CI [.06, .22], *p* < .001. However, older adults did not significantly differ in their curiosity for low-upvote items across the upvote visible condition (*M* = .54, SE = .03) and upvote invisible condition (*M* = .58, SE = .03), *b_estimate_* = .05, SE = .04, 95% CI [-.24, .13], *p* = .178. There were no differences in younger and older adults’ curiosity ratings for high-upvote items across conditions (all *p*s > .05).

**Fig 4.**
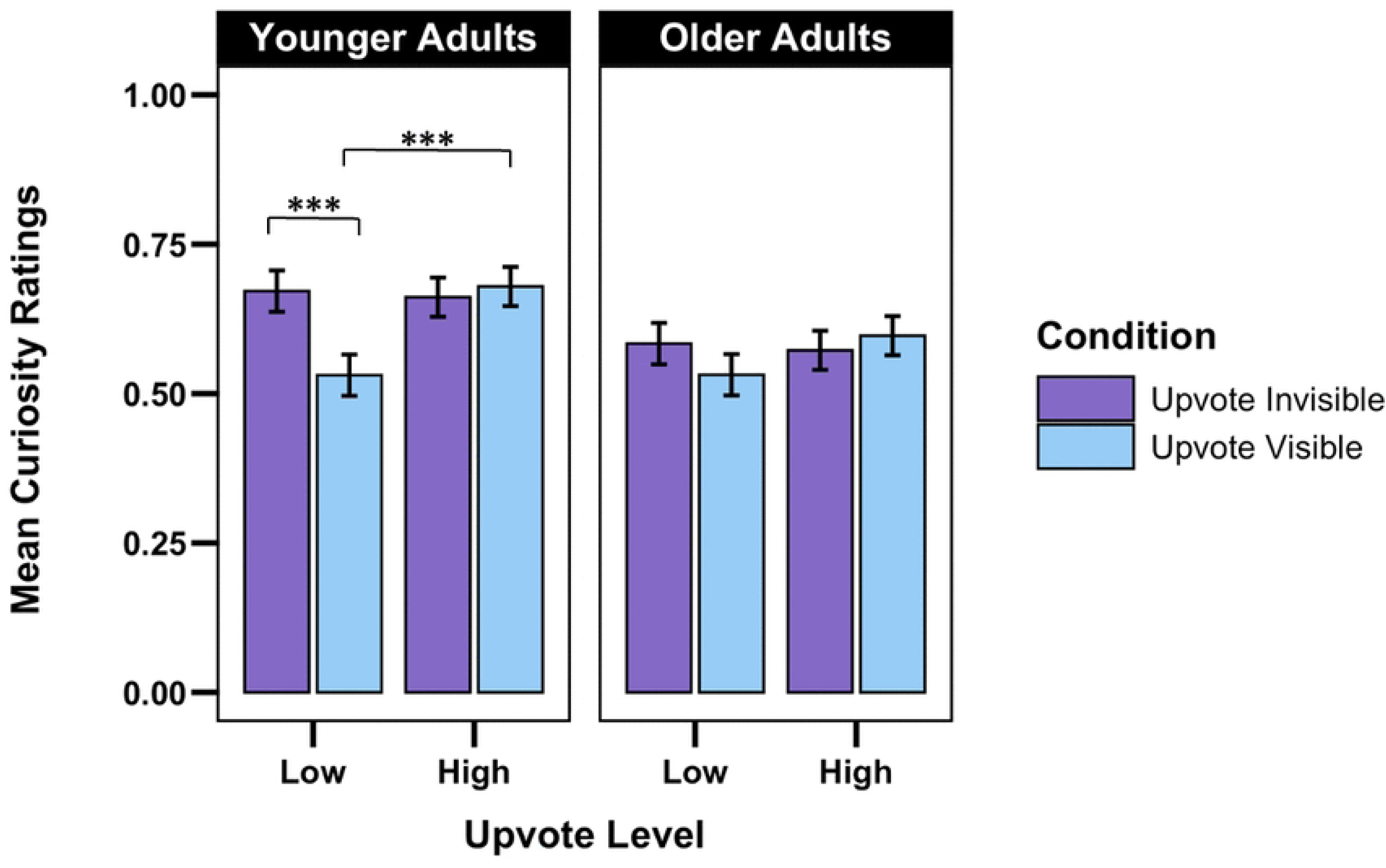
Curiosity Ratings in Experiment 2. \*\*\**p* < .001. Error bars represent one standard error above and below the estimated marginal mean.

Parsed by Condition and Upvote Level, younger adults were more curious about high- upvote items (*M* = .68, SE = .03) than older adults (*M* = .60, SE = .03) in the upvote visible condition, *b*_estimate_ = -.08, SE = .04, 95% CI [-.15, -.01], *p* = .019. However, there was no difference between younger (*M* = .53, SE = .03) and older adults (*M* = .53, SE = .03) in their curiosity for low-upvote items in the upvote visible condition, *b*_estimate_ = .00, SE = .04, 95% CI [-.08, .08], *p* = .987. Moreover, in the upvote invisible condition, younger adults were more curious about high-upvote items (*M* = .66, SE = .03) than older adults (*M* = .57, SE = .03), *b*_estimate_ = -.09, SE = .04, 95% CI [-.16, -.02], *p* = .011. Similarly, in the upvote invisible condition, younger adults were more curious about low-upvote items (*M* = .67, SE = .03) than older adults (*M* = .58, SE = .03), *b*_estimate_ = -.09, SE = .04, 95% CI [-.16, -.01], *p* = .024.

Parsed by Age and Condition, in the upvote visible condition, younger adults were more curious about high- (*M* = .68, SE = .03) than low-upvote items (*M* = .53, SE = .03), *b*_estimate_ = .15, SE = .04, 95% CI [.08, .22], *p* < .001. There was no significant difference between older adults’ curiosity for high- and low-upvote items in the upvote visible condition, or younger or older adults’ curiosity about high- compared to low-upvote items in the upvote invisible condition (all *p*s > .05).

In summary, curiosity ratings were sensitive to social value cues, although this sensitivity varied by age and condition. Younger adults were more curious about high- than low-upvote items in the upvote visible condition, whereas older adults did not show a significant difference. Neither age group showed significant differences in curiosity between high- and low-upvote items in the upvote invisible condition. Additionally, younger adults were less curious about low-upvote items in the upvote visible than in the upvote invisible condition, whereas older adults did not differ significantly across conditions. Notably, younger adults were more curious than older adults about both high- and low-upvote items in the upvote invisible condition, as well as high-upvote items in the upvote visible condition. These findings suggest that younger adults’ curiosity was more responsive to social value cues than that of older adults, while age-related differences in curiosity were also evident in the absence of visible social information.

### Recall performance

To identify the best-fitting model, we conducted a series of nested model comparisons using likelihood ratio tests. The final model that converged included a random by-participant and by-question intercept, excluding all correlations among random effects.

We found a significant main effect of age, *χ^2^* (1) = 12.85, *p* < .001 (Fig 5), such that semantic similarity scores were higher for older adults (*M* = .40, SE = .02) than for younger adults (*M* = .32, SE = .27). No other effects were significant (all *ps >* .05), suggesting that social value cues did not significantly influence recall of trivia answers in either age group.

**Fig 5.**
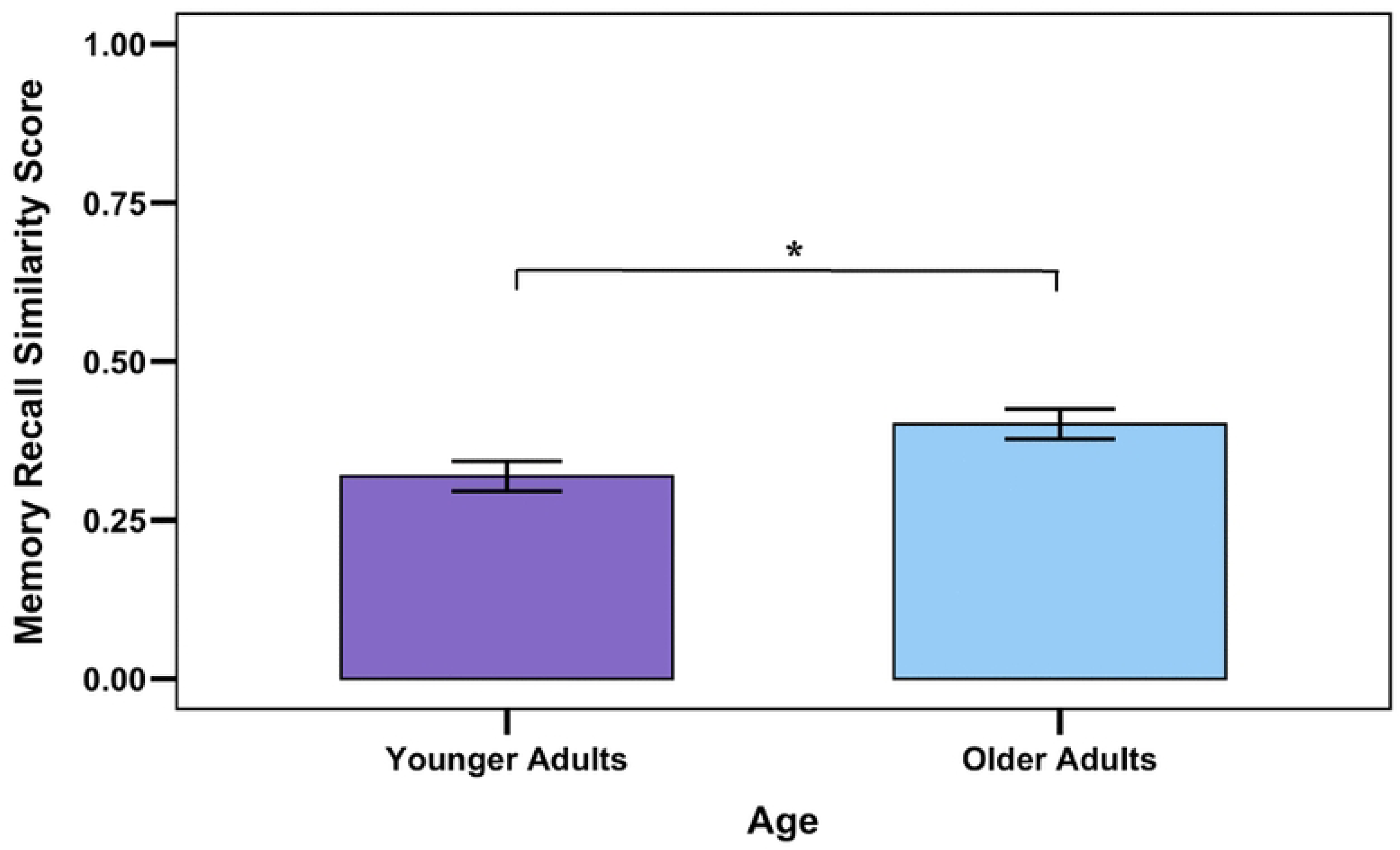
Recall Accuracy by Age Group in Experiment 2. \**p* < .05.

## Discussion

The current study examined how social value cues affect curiosity, information seeking, and memory in younger and older adults. We aimed to replicate previous work [8] that had found age differences in social value effects on curiosity, and in curiosity effects on information seeking. Crucially, we also sought to test whether the impact of social value cues would extend to subsequent trivia recall – an important question given the functional importance of episodic memory across the lifespan. In both experiments, we manipulated the perceived social value of trivia questions through upvotes, a popularity metric used on social media platforms. Results showed that social value cues influenced perceptions of popularity as well as curiosity in both younger and older adults. However, social value cues did not have a significant impact on information-seeking behavior or on memory performance. We discuss each of these findings in turn.

## Effects of social value cues on perceived popularity

In both experiments, participants in both age groups rated high-upvote items as being more popular than low-upvote items when upvotes were visible. However, popularity ratings were influenced more by upvotes in younger adults than in older adults. It is possible that upvotes were less effective for older adults because older adults have less experience with these metrics. It should be noted, however, that we tested participants’ understanding of upvotes during a comprehension check before the experiment. Another possibility is that older adults may have read the questions more closely than younger adults, and that older adults’ popularity ratings may thus have been more strongly influenced by the content of the questions and less strongly by upvotes. This interpretation would fit with the observation that older adults are more highly motivated than younger adults to achieve high accuracy in psychological studies [46].

## Effects of social value on curiosity

In both experiments, social value cues affected curiosity ratings in both age groups, as expressed in significant Condition x Upvote Level interactions. In the first experiment, we found higher curiosity for high-upvote vs. low-upvote items in the upvote visible condition, but not in the upvote invisible condition where upvotes were not visible.

In Experiment 2, this pattern was more nuanced, with the effects of social value cues varying by age. Specifically, younger adults were more curious about high- than low-upvote items in the upvote visible condition, whereas older adults did not show a significant difference. Additionally, younger adults were less curious about low-upvote items in the upvote visible than in the upvote invisible condition, whereas older adults did not differ significantly across conditions. Although these findings indicate that curiosity is sensitive to social value cues across the adult lifespan, our predictions regarding age differences were only partially supported. Based on Swirsky and Spaniol (2024, Exp. 1), we had expected to find an age-related positivity effect in social value effects, such that older adults would be more curious about high-upvote (“positive”) items in the upvote visible compared with the upvote invisible condition, whereas younger adults would be less curious about low-upvote (“negative”) items in the upvote visible compared to the upvote invisible condition. Contrary to this hypothesis, in Experiment 1, age did not significantly moderate the social-value effect. In Experiment 2, while younger adults showed less curiosity for low-upvote items when upvotes were visible compared to invisible, there was no difference in older adults’ curiosity for high-upvote items across conditions. Moreover, in the latter study, younger adults were more curious than older adults about both high- and low-upvote items when upvotes were invisible, suggesting that age-related differences in curiosity were not exclusively attributable to social value cues.

What may have caused this inconsistent pattern of age differences? As noted, Swirsky & Spaniol’s (2024, Exp. 1) methodology had failed to disentangle social cue effects from item effects (i.e., high-upvote items were genuinely more popular than low-upvote items). To control for this confound, the current experiments used only medium-popularity items and artificially assigned them high or low upvotes. Nevertheless, the age differences observed in Experiment 2 in the upvote invisible condition suggest that factors beyond experimentally manipulated social value cues may have contributed to curiosity ratings. It is possible that age differences in social cue effects on curiosity depend on the exact nature of the information that is being probed, or on the congruency between items and social cues. For example, what is considered positive information under SST likely extends beyond social value, and may depend on the content of the question, such as reference to death in one of the items, “How can alcohol withdrawal or detox kill you?” Moreover, certain questions may be of increased personal relevance to a given age group, impacting curiosity. Using the example of alcohol withdrawal, this question may be of greater curiosity to younger adults, who are more impacted by alcohol use disorders than older adults [47]. The use of basic science questions that are emotionally neutral and less pertinent to a given age group (e.g., “How do scientists know what the global temperature was millions of years ago?”) may prove optimal in this regard. While this issue remains to be clarified in future research, the current findings provide evidence that social value cues influence curiosity in both younger and older adults, although the magnitude of these effects may vary by age and experimental context. Importantly, the age differences observed in Experiment 2 in the absence of visible social value cues suggest that curiosity is also shaped by factors beyond social endorsement. Nevertheless, our findings support the notion that other people’s endorsement (or lack thereof) can influence curiosity [22].

## Effects of social value on information seeking

Contrary to our predictions, information seeking was not influenced by social value cues in Experiment 1. This null finding aligns with Swirsky and Spaniol (2024), who also found no systematic variation in information seeking based on social value. Notably, this absence of effects on answer reveal occurred despite the curiosity ratings showing sensitivity to social value cues. One interpretation of these findings is that curiosity ratings and answer reveal behavior reflect partially distinct processes. Social value cues may influence an initial appraisal of an item as interesting or worthy of attention, but the decision to invest effort in obtaining the answer may depend more strongly on intrinsic factors. These factors may include prior knowledge, perceived usefulness of the information, and the effort required to obtain it [13,20,21,23,48]. For example, a participant may report elevated curiosity when a question appears highly endorsed by others, yet still choose not to reveal the answer if the topic is personally irrelevant or if the expected informational value is low. Thus, social endorsement may be sufficient to shape subjective curiosity, but insufficient to consistently drive information-seeking behavior.

## Effects of social value on memory

Lastly, the results of Experiment 2 showed that social value cues had no significant impact on memory for information, in either age group.

Similar to information seeking, accurate memory retrieval may be determined by a broader set of factors than initial curiosity alone. Specifically, recall may also be influenced by prior knowledge of the topic, pre-existing personal interest, attentional engagement during encoding, and the depth of processing applied to the information [3,49–51]. These intrinsic factors may have exerted a stronger influence on memory performance than the social value cues.

Interestingly, recall accuracy was higher in older adults than in younger adults – a finding that contradicts typical patterns of age-related decline in episodic memory tasks [52]. Older adults, on average, provided slightly longer responses (*M =* 15.08 words) compared to younger adults (*M* = 12.88 words). While this may signal greater effort overall from older adults, consistent with evidence that they are often more motivated to perform accurately in research settings (46), a t-test evaluating this relationship was nonsignificant, *t*(172.95) =1.22, *p* = .112, suggesting that operationalizations of effort in terms of verbosity cannot explain higher recall accuracy in older adults than younger adults. Alternatively, the trivia answers were relatively long and semantically complex, which may have increased encoding demands and reduced the likelihood that a modest increase in curiosity would translate into measurable memory benefits [53].

Finally, we cannot rule out that some participants may have relied on generative artificial intelligence or internet searches to generate their answers, rather than retrieving the information from memory. In a few instances (n=2), we observed that different participants provided identical or nearly identical responses to the same question, including additional details that were not present in the original answer. While the removal of these two data points did not affect the results, these patterns suggest that participants may have conducted a search of the information during the recall phase rather than relying on their memory. If such behavior occurred, it could have introduced noise into the memory data and obscured potential effects of social value cues.

## Strengths, limitations, and future directions

The present study has several strengths and limitations that can guide future research. First, the study clarifies previous findings regarding the effects of social value cues on curiosity and information seeking. By comparing an upvote-visible condition with an upvote invisible control condition while controlling item assignments, the present work suggests that previously reported effects may have been influenced, at least in part, by differences in the intrinsic appeal of trivia items rather than social endorsement alone. The study also benefited from a preregistered design and a relatively large sample of younger and older adults recruited from two online participant pools, enhancing transparency and generalizability.

Another strength is the novel use of transformer-based language models to evaluate recall accuracy [39,43]. Traditional scoring methods for free recall are often binary or manual, which is labour-intensive and may overlook nuance. In contrast, transformer-based language models offer scalable methods to index free recall, whereby continuous semantic similarity scores capture overlap in meaning between participants’ responses and the correct answers [41]. This property makes transformer-based language models particularly useful for trivia tasks, where recall may be partial or paraphrased. By incorporating a machine learning-based method, this method not only improves scoring reproducibility and objectivity but also sets a precedent for future curiosity and memory research.

A limitation relates to the use of trivia to evaluate memory. In using real trivia questions in our memory paradigm, we may have captured both episodic memory for the studied information and participants’ pre-existing semantic knowledge. To overcome this limitation, it would be useful, in future work, to examine the impact of prior knowledge on curiosity and subsequent memory more directly, for example by tapping into participants’ idiosyncratic areas of expertise.

Finally, the use of fixed trivia items paired with manipulated upvotes is another limitation of the present study. Although item-upvote assignments were randomized and the trivia questions were selected to have moderate baseline popularity, intrinsic differences in item interest were likely not completely eliminated. As a result, some effects may have reflected residual differences in item appeal rather than experimentally manipulated social value. Future work should continue developing paradigms that more effectively separate the influence of social endorsement from the inherent characteristics of the information itself.

## Conclusion

The present study investigated how social value cues, operationalized via upvote levels and their visibility, affect curiosity, information seeking, and recall in younger and older adults. Despite successfully manipulating perceptions of popularity, effects on curiosity, information search, and memory were nuanced, suggesting that social cues may play a limited role in motivating curiosity-driven behaviour. Instead, internal factors such as effort and individual engagement may have exerted a stronger influence. Methodologically, this study introduced computationally derived semantic similarity as a novel way to assess recall in curiosity research, since it can capture accurate recall via shared meaning or concepts rather than exact word matching. While this represents a key strength of the study, some limitations include the use of trivia items to test memory and limited control over item-level interest. Overall, these findings suggest that while social endorsement may shape perceptions of informational value, its ability to drive curiosity, exploration, or memory may be constrained when other motivational factors are involved. Importantly, this study advances understanding of curiosity and motivation-cognition interactions by highlighting how internal and external influences interact in younger and older adults. Future research should aim to disentangle these influences further by incorporating greater control over content relevance, participant effort, and internal motivational states.

## Acknowledgements

The authors sincerely thank the anonymous peer reviewers for their thoughtful manuscript reviews and constructive feedback, which significantly improved the clarity of this paper. We would also like to thank Drs. Karl Szpunar and Todd Girard for their invaluable guidance, insights, and feedback while serving on the thesis committee for this project. Additionally, we extend our gratitude to Dr. Lindsay Santacroce for generously sharing her technical expertise with PsychoPy and Pavlovia, which was instrumental to our experimental setup.

## Supporting information

S1 Fig 1. Popularity Ratings by Condition and Upvote Level Across Age Groups (Experiment 1). \*\**p* < .01, \*\*\**p* < .001.

S1 Fig 2. Popularity Ratings by Condition and Upvote Level Across Age Groups (Experiment 2). \*\*\**p* < .001.

## Notes

### Competing Interest Statement

The authors have declared no competing interest.

## References

1. Chu L, Fung HH. Age Differences in State Curiosity: Examining the Role of Personal Relevance. Gerontology. 2022;68(3):321–9. doi:10.1159/000516296

2. Galli G, Sirota M, Gruber MJ, Ivanof BE, Ganesh J, Materassi M, et al. Learning facts during aging: the benefits of curiosity. Experimental Aging Research. 2018 Aug 8;44(4):311–28. doi:10.1080/0361073X.2018.1477355

3. McGillivray S, Murayama K, Castel AD. Thirst for knowledge: The effects of curiosity and interest on memory in younger and older adults. Psychology and Aging. 2015 Dec;30(4):835–41. doi:10.1037/a0039801

4. Robinson OC, Demetre JD, Litman JA. Adult life stage and crisis as predictors of curiosity and authenticity: Testing inferences from Erikson’s lifespan theory. International Journal of Behavioral Development. 2017 May;41(3):426–31. doi:10.1177/0165025416645201

5. Litman JA, Silvia PJ. The Latent Structure of Trait Curiosity: Evidence for Interest and Deprivation Curiosity Dimensions. Journal of Personality Assessment. 2006 Jun;86(3):318–28. doi:10.1207/s15327752jpa8603_07

6. Whatley MC, Murayama K, Sakaki M, Castel AD. Curiosity across the adult lifespan: Age- related differences in state and trait curiosity. Zhu G, editor. PLoS ONE. 2025 May 7;20(5):e0320600. doi:10.1371/journal.pone.0320600

7. Sakaki M, Yagi A, Murayama K. Curiosity in old age: A possible key to achieving adaptive aging. Neuroscience & Biobehavioral Reviews. 2018 May;88:106–16. doi:10.1016/j.neubiorev.2018.03.007

8. Swirsky LT, Spaniol J. Moderators of curiosity and information seeking in younger and older adults. Psychology and Aging. 2024 Nov;39(7):701–14. doi:10.1037/pag0000847

9. Gruber MJ, Ranganath C. How Curiosity Enhances Hippocampus-Dependent Memory: The Prediction, Appraisal, Curiosity, and Exploration (PACE) Framework. Trends in Cognitive Sciences. 2019 Dec;23(12):1014–25. doi:10.1016/j.tics.2019.10.003

10. Duarte A, Kensinger E. Age-related changes in episodic memory. In: Samanez-Larkin GR, editor. The aging brain: Functional adaptation across adulthood. [Internet]. Washington: American Psychological Association; 2019 [cited 2026 Sep 3]. p. 111-34. Available from: https://content.apa.org/books/16123-005 doi:10.1037/0000143-005

11. Swirsky LT, Shulman A, Spaniol J. The interaction of curiosity and reward on long-term memory in younger and older adults. Psychology and Aging. 2021 Aug;36(5):584–603. doi:10.1037/pag0000623

12. Swirsky LT, Spaniol J. Consequences of curiosity for recognition memory in younger and older adults. Psychon Bull Rev. 2023 Dec 14. doi:10.3758/s13423-023-02414-y

13. Gottlieb J, Oudeyer PY, Lopes M, Baranes A. Information-seeking, curiosity, and attention: computational and neural mechanisms. Trends in Cognitive Sciences. 2013 Nov;17(11):585–93. doi:10.1016/j.tics.2013.09.001

14. Kelly ChristopherA, Sharot T. Individual differences in information-seeking. Nat Commun. 2021 Dec 3;12(1):7062. doi:10.1038/s41467-021-27046-5

15. Spreng RN, Turner GR. From exploration to exploitation: a shifting mental mode in late life development. Trends in Cognitive Sciences. 2021 Dec;25(12):1058–71. doi:10.1016/j.tics.2021.09.001

16. Oudeyer PY, Kaplan F. What is intrinsic motivation? A typology of computational approaches. Front Neurorobot. 2007;1. doi:10.3389/neuro.12.006.2007

17. Wade S, Kidd C. The role of prior knowledge and curiosity in learning. Psychon Bull Rev. 2019 Aug;26(4):1377–87. doi:10.3758/s13423-019-01598-6

18. Loewenstein G. The psychology of curiosity: A review and reinterpretation. Psychological Bulletin. 1994 Jul;116(1):75–98. doi:10.1037/0033-2909.116.1.75

19. Metcalfe J, Schwartz BL, Eich TS. Epistemic curiosity and the region of proximal learning. Current Opinion in Behavioral Sciences. 2020 Oct;35:40–7. doi:10.1016/j.cobeha.2020.06.007

20. Metcalfe J, Kornell N. A Region of Proximal Learning model of study time allocation. Journal of Memory and Language. 2005 May;52(4):463–77. doi:10.1016/j.jml.2004.12.001

21. Dubey R, Griffiths TL. Reconciling novelty and complexity through a rational analysis of curiosity. Psychological Review. 2020 Apr;127(3):455–76. doi:10.1037/rev0000175

22. Dubey R, Mehta H, Lombrozo T. Curiosity Is Contagious: A Social Influence Intervention to Induce Curiosity. Cognitive Science. 2021 Feb;45(2):e12937. doi:10.1111/cogs.12937

23. Dubey R, Griffiths TL, Lombrozo T. If it’s important, then I’m curious: Increasing perceived usefulness stimulates curiosity. Cognition. 2022 Sep;226:105193. doi:10.1016/j.cognition.2022.105193

24. Carstensen LL, Isaacowitz DM, Charles ST. Taking time seriously: A theory of socioemotional selectivity. American Psychologist. 1999;54(3):165–81. doi:10.1037/0003-066X.54.3.165

25. Mather M, Carstensen LL. Aging and motivated cognition: the positivity effect in attention and memory. Trends in Cognitive Sciences. 2005 Oct;9(10):496–502. doi:10.1016/j.tics.2005.08.005

26. Campbell JID, Thompson VA. MorePower 6.0 for ANOVA with relational confidence intervals and Bayesian analysis. Behav Res. 2012 Dec;44(4):1255–65. doi:10.3758/s13428-012-0186-0

27. Litman JA, Collins RP, Spielberger CD. The nature and measurement of sensory curiosity. Personality and Individual Differences. 2005 Oct;39(6):1123–33. doi:10.1016/j.paid.2005.05.001

28. Collins RP, Litman JA, Spielberger CD. The measurement of perceptual curiosity. Personality and Individual Differences. 2004 Apr;36(5):1127–41. doi:10.1016/S0191-8869(03)00205-8

29. Raven JC. Guide to using the Mill Hill Vocabulary Scale with the Progressive Matrices Scales. H. K. Lewis & Co.; 1958.

30. Watson D, Clark LA, Tellegen A. Development and validation of brief measures of positive and negative affect: The PANAS scales. Journal of Personality and Social Psychology. 1988;54(6):1063–70. doi:10.1037/0022-3514.54.6.1063

31. Lovibond PF, Lovibond SH. The structure of negative emotional states: Comparison of the Depression Anxiety Stress Scales (DASS) with the Beck Depression and Anxiety Inventories. Behaviour Research and Therapy. 1995 Mar;33(3):335–43. doi:10.1016/0005-7967(94)00075-U

32. Peirce J, Gray JR, Simpson S, MacAskill M, Höchenberger R, Sogo H, et al. PsychoPy2: Experiments in behavior made easy. Behav Res. 2019 Feb;51(1):195–203. doi:10.3758/s13428-018-01193-y

33. Brown VA. An Introduction to Linear Mixed-Effects Modeling in R. Advances in Methods and Practices in Psychological Science. 2021 Jan;4(1):2515245920960351. doi:10.1177/2515245920960351

34. Bates D, Mächler M, Bolker B, Walker S. Fitting linear mixed-effects models using lme4. Vol. J67. 2015;J67(1):1-48. 10.18637/jss.v067.i01

35. Murayama K, Sakaki M, Yan VX, Smith GM. Type I error inflation in the traditional by- participant analysis to metamemory accuracy: A generalized mixed-effects model perspective. Journal of Experimental Psychology: Learning, Memory, and Cognition. 2014 Sep;40(5):1287–306. doi:10.1037/a0036914

36. Szumilas M. Explaining odds ratios. J Can Acad Child Adolesc Psychiatry. 2010 Aug;19(3):227–9. PubMed PMID: 20842279; PubMed Central PMCID: PMC2938757.

37. Fox J, Weisberg S. An R companion to applied regression. Thousand Oaks: Sage Publishing; 2011.

38. Lenth R, Singmann H, Love J, Buerkner P, Herve M. Package “Emmeans”. [Internet]. 2018. Available from: http://cran.r-project.org/package=emmeans

39. Shen X, Houser T, Smith DV, Murty VP. Machine-learning as a validated tool to characterize individual differences in free recall of naturalistic events. Psychon Bull Rev. 2023 Feb;30(1):308–16. doi:10.3758/s13423-022-02171-4

40. Wang J, Dong Y. Measurement of Text Similarity: A Survey. Information. 2020 Aug 31;11(9):421. doi:10.3390/info11090421

41. Devlin J, Chang MW, Lee K, Toutanova K. BERT: Pre-training of deep bidirectional transformers for language understanding. In: Proceedings of the 2019 Conference of the North American Chapter of the Association for Computational Linguistics: Human Language Technologies, Volume 1 (Long and Short Papers) [Internet]. Minneapolis, Minnesota: Association for Computational Linguistics; 2019 [cited 2026 Sep 3]. p. 4171-86. Available from: https://aclanthology.org/N19-1423 doi:10.18653/v1/N19-1423

42. Mikolov T, Chen K, Corrado G, Dean J. Efficient estimation of word representations in vector space [Internet]. arXiv; 2013 [cited 2026 Sep 3]. Available from: https://arxiv.org/abs/1301.3781 doi:10.48550/ARXIV.1301.3781

43. Fowler Z, Palombo DJ, Madan CR, O’Connor BB. Collaborative imagination synchronizes representations of the future and fosters social connection in the present. Proc Natl Acad Sci USA. 2024 Jun 18;121(25):e2318292121. doi:10.1073/pnas.2318292121

44. Reimers N, Gurevych I. Sentence-BERT: Sentence Embeddings using Siamese BERT- Networks. In: Proceedings of the 2019 Conference on Empirical Methods in Natural Language Processing and the 9th International Joint Conference on Natural Language Processing (EMNLP-IJCNLP) [Internet]. Hong Kong, China: Association for Computational Linguistics; 2019 [cited 2026 Sep 14]. p. 3980-90. Available from: https://www.aclweb.org/anthology/D19-1410 doi:10.18653/v1/D19-1410

45. Sanh V, Debut L, Chaumond J, Wolf T. DistilBERT, a distilled version of BERT: smaller, faster, cheaper and lighter [Internet]. arXiv; 2019 [cited 2026 Sep 3]. Available from: https://arxiv.org/abs/1910.01108 doi:10.48550/ARXIV.1910.01108

46. Ryan AD, Campbell KL. The ironic effect of older adults’ increased task motivation: Implications for neurocognitive aging. Psychon Bull Rev. 2021 Dec;28(6):1743–54. doi:10.3758/s13423-021-01963-4

47. Veerbeek MA, Ten Have M, Van Dorsselaer SA, Oude Voshaar RC, Rhebergen D, Willemse BM. Differences in alcohol use between younger and older people: Results from a general population study. Drug and Alcohol Dependence. 2019 Sep;202:18–23. doi:10.1016/j.drugalcdep.2019.04.023

48. Payne K. The impact of effort on the search for information. Kansas State University; 2025.

49. Brod G, Werkle-Bergner M, Shing YL. The influence of prior knowledge on memory: A developmental cognitive neuroscience perspective. Front Behav Neurosci. 2013;7. doi:10.3389/fnbeh.2013.00139

50. Cowan N, Bao C, Bishop-Chrzanowski BM, Costa AN, Greene NR, Guitard D, et al. The relation between attention and memory. Annu Rev Psychol. 2024 Jan 18;75(1):183–214. doi:10.1146/annurev-psych-040723-012736

51. Craik FIM, Lockhart RS. Levels of processing: A framework for memory research. Journal of Verbal Learning and Verbal Behavior. 1972 Dec;11(6):671–84. doi:10.1016/S0022-5371(72)80001-X

52. Cansino S. Episodic memory decay along the adult lifespan: A review of behavioral and neurophysiological evidence. International Journal of Psychophysiology. 2009 Jan;71(1):64–9. doi:10.1016/j.ijpsycho.2008.07.005

53. Rothkopf EZ, Billington MJ. Passage length and recall with test size held constant: effects of modality, pacing, and learning set. Journal of Verbal Learning and Verbal Behavior. 1983 Dec;22(6):667–81. doi:10.1016/S0022-5371(83)90395-X

